# A prior exposure to *Serratia marcescens* or xenobiotics primes *Drosophila* enterocytes against a recurring cytoplasmic purge

**DOI:** 10.1101/2021.10.31.466690

**Authors:** Simone Terreri, Bechara Sina-Rahme, Inês Pais, Catherine Socha, Matthieu Lestradet, Miriam Yamba, Stefanie Schirmeier, Kwang-Zin Lee, Dominique Ferrandon

## Abstract

The cytoplasmic extrusion of enterocytes is a fast response to an exposure to pore-forming toxin (PFT)-producing bacteria whereby their apical cytoplasm is extruded into the intestinal lumen. As a result of this purge, the intestinal epithelium becomes thin prior to a subsequent recovery. We report here that the ingestion of ethanol or caffeine induces a similar response, which suggests that a common purging process is triggered by bacterial toxins and abiotic toxicants. We also delineate an additional mechanism that is initiated by these stimuli that we refer to as priming. The initial exposure of the intestinal epithelium to either PFT or xenobiotics protects enterocytes against a further round of purging upon a second bacterial infection. Priming prevents the epithelium from being persistently thin in the context of chronic intestinal infections. We have identified the upper part of the p38b MAPK pathway as well as the homeobox-containing transcription factors E5/EMS as being required for priming and not for the regrowth of enterocytes after the cytoplasmic purge. Unexpectedly, the priming process appears to function cell-nonautonomously. Our findings suggest that the cytoplasmic purge extrusion has been selected because it constitutes a fast reaction to accidental exposure to bacterial toxins or toxicants.

## Introduction

*Drosophila* flies preferentially feed on decaying fruits, which constitute the stage of an ongoing microbial warfare. For instance, yeast such as *Saccharomyces* limit microbial competition by releasing a toxicant effective against bacteria, ethanol. Interestingly, whereas most insects lack an ability to tolerate ethanol, *Drosophila* have a functional alcohol dehydrogenase enzyme (Ashburner, 1998). Nowadays, insects are also exposed to many xenobiotics that may lace their food. The gut epithelium constitutes a major frontier between the organism and the food it ingests. It must be able to digest nutrients while protecting its host from the noxious effects of ingested pathogens or chemicals that may occasionally contaminate its food.

Whereas many studies have documented the effects of the ingestion of pathogens or selected xenobiotics on the homeostasis of the intestinal epithelium from the perspective of intestinal stem cell (ISC) compensatory proliferation to replace damaged enterocytes, additional regulatory mechanisms may be at play such as the growth of enterocytes (Bonfini et al., 2021; Bonfini et al., 2016; Lemaitre and Miguel-Aliaga, 2013; O’Brien et al., 2011; Tamamouna et al., 2020; Xiang et al., 2017; Zhang and Edgar, 2021). One original mechanism we have reported in the framework of our studies on host defenses against ingested pathogens such as *Serratia marcescens* (strain Db11: *SmDb11*) is that of the cytoplasmic extrusion of enterocytes (Lee et al., 2016). In response to the exposure to pore-forming toxins, several events have been documented. First, there is a transient formation of lipid droplets in the apical part of enterocytes, which form prominent dome-like structures in the R2 region of the gut (Buchon et al., 2013; Shanbhag and Tripathi, 2005). Mitochondria fuse in the apical cytoplasm and form megamitochondria that are likely not functional. The next event consists in the extrusion of this apical cytoplasm through large apertures in the apical cytoplasmic membrane that release and eliminate the apical cytoplasm that contains damaged organelles and likely the pore-forming toxins as well as invading bacteria. As a result, a thin epithelium is observed within one to three hours of the ingestion of *SmDb11*. This thin epithelium is not the consequence of the spreading of enterocytes but of an important loss of their volume. This cytoplasmic purge may alleviate the stress exerted by pathogens on enterocytes. Indeed, we failed to observe an induction of stress-response pathway such as the Jun Kinase (JNK) or the Janus-Kinase/ Signal Transducer and Activator of Transcription (JAK-STAT) pathways. We did not observe an increase in necrosis or apoptosis nor an increase in the proliferation rate of ISCs (Lee et al., 2016).

The thin epithelium stage is followed by a rapid recovery of the original shape and volume of the enterocytes in the following hours. This restoration occurs within 12 to 24 hours, even though organelles such as the endoplasmic reticulum still appear to be somewhat disorganized. We have identified *Cyclin J* (*CycJ*) as a gene likely required for this recovery process since the intestinal epithelium of *CycJ* mutant flies remains thin upon chronic *SmDb11* oral infection. The phenomenon of the apical cytoplasmic purge constitutes an evolutionarily conserved fast response to the exposure to pore-forming toxins that represents a novel resilience mechanism (Lee et al., 2016).

Here, we first address the question as to whether the cytoplasmic purge might also be at play upon the ingestion of xenobiotics that the fly may (ethanol) or may not (caffeine) encounter in its current natural environment. We next report the existence of a priming mechanism that prevents the intestinal epithelium from being persistently thin during chronic gut infections. Remarkably, the ingestion of xenobiotics provides cross-priming protection against a secondary challenge with *SmDb11*. Finally, we have identified genes required for this priming phenomenon that unexpectedly function noncell-autonomously.

## Results

### The ingestion of caffeine or ethanol leads to the cytoplasmic extrusion of the apical part of enterocytes

We have exposed flies either to caffeine, a strong insecticide or to ethanol, a xenobiotic well tolerated by *D. melanogaster*. As in our prior study, we have focused on the R2 region of the anterior midgut (Buchon et al., 2013; Lee et al., 2016). We usually observed a rapid thinning of the intestinal epithelium following the ingestion (Fig.1A). It appeared however to be faster and shorter-lived as that induced by exposure to *SmDb11* (Fig. 1B-C) and often the size of neighboring enterocytes did not appear to be as homogenous as that observed upon a challenge with hemolysin. The extent of thinning appeared also to be not as extensive (Fig. 1). Nevertheless, the cytoplasmic purge induced by xenobiotics exhibited features observed using *SmDb11* such as the formation of lipid droplets in enterocytes (Fig. 2A-C) and of megamitochondria (Fig. 2D) in the apical part of the enterocytes. The next step was to determine whether thinning correlated with a cytoplasmic purge. To visualize the cytoplasm of enterocytes, we expressed GFP in their cytoplasm and indeed observed the extrusion of the labelled cytoplasm of enterocytes into the ectoperitrophic space of the gut lumen (Fig. 2E). In keeping with this observation, we observed the extrusion of lipid droplets as well as that of megamitochondria (Fig. 2B-C, F, Fig. S1 B-C). As the formation of megamitochondria may result from oxidative stress, we monitored the redox state of the cytoplasm and mitochondria using dedicated ratiometric reporters (Albrecht et al., 2011; Shutt et al., 2012). We found that only caffeine significantly increased the oxidation state in mitochondria whereas we did not observe any modification of the redox potential in the cytoplasm (Fig. S1A). Of note, megamitochondria appeared to more reliably detected after caffeine challenge than after the ingestion of ethanol. We have also checked in survival experiments that caffeine rapidly kills the fly. In contrast, we observed a mild loss of fitness of flies exposed to ethanol only when it was provided everyday (Fig. S1D-E). We also indirectly monitored whether any damage was inflicted to enterocytes by measuring the rate of division of ISCs and observed a significant enhanced proliferation after the ingestion of *SmDb11*, as previously reported, and of ethanol or paraquat (Fig. S1F). Interestingly, caffeine did not induce any compensatory proliferation of ISCs, a phenomenon that will require further study. Paraquat is highly toxic and was used here as a positive control. We have tested also other xenobiotics for a possible purge including paraquat, a heavy metal (Cd), and an inhibitor of the mitochondrial respiratory chain, rotenone. We have observed the formation of lipid droplets and megamitochondria after the ingestion of metals or paraquat and also some epithelial thinning also occurring after this challenge (Fig. S1G-H). The situation was complex however because of the toxicity of these compounds. For instance, we detected the formation of megamitochondria as early as one hour after ingestion of paraquat but a very thin epithelium only some seven hours after the ingestion of paraquat, which was nevertheless followed by a recovery of the original thickness (Fig. S1H).

**Figure 1:**
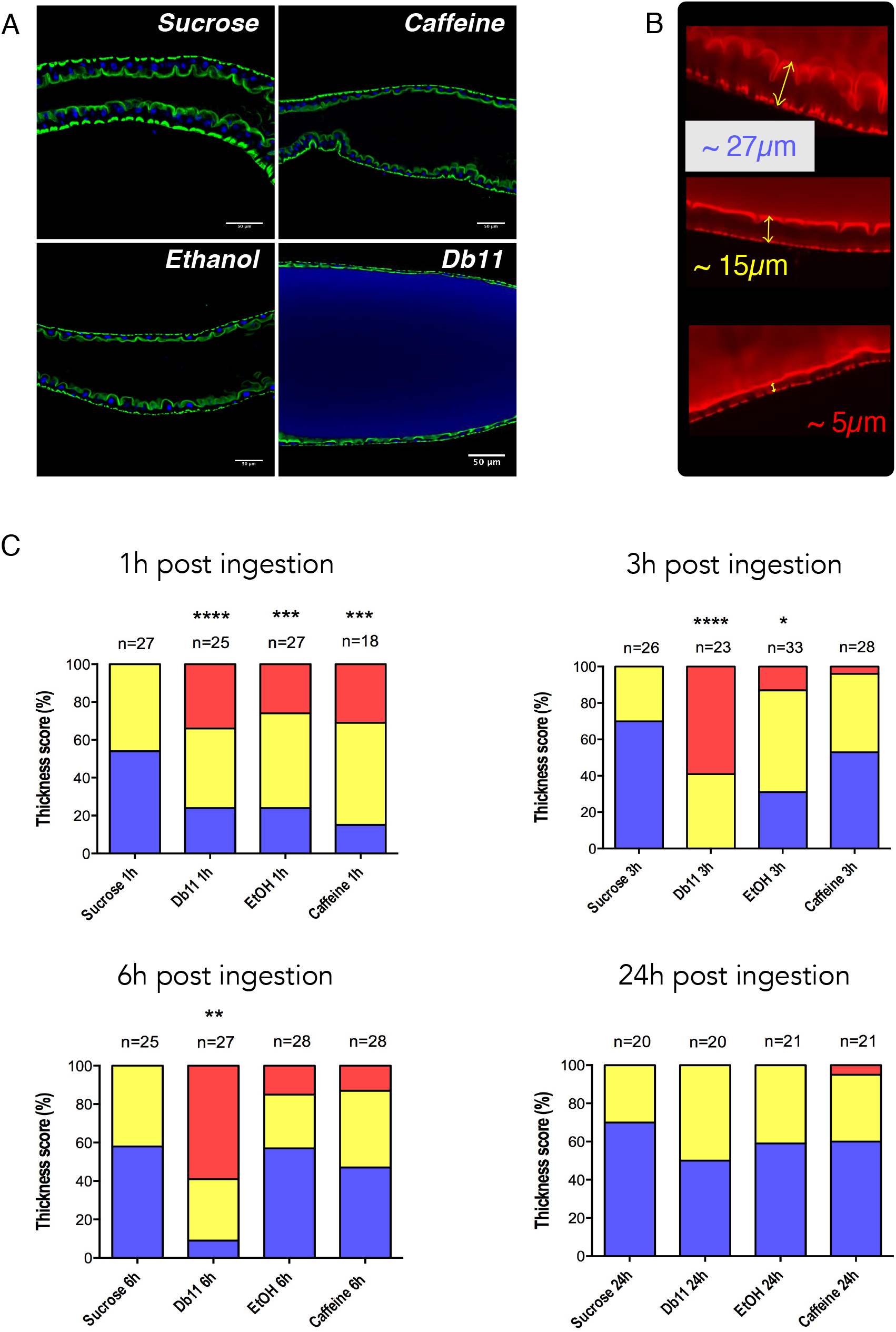
Ethanol and caffeine also induce epithelial thinning. **(A)** Confocal pictures of midguts of *w*^*A5001*^ flies fed with sucrose 100mM, Ethanol 2,5%, Caffeine 2,5mg/mL, or Db11 OD_600_=10. Flies were dissected 1h after ingestion, unlike Db11 dissected 3h post ingestion. Scale bar: 50μm. **(B)** Confocal pictures of the different classes of midgut thicknesses used for the qualitative scoring of observed guts from normal, intermediate and reduced thickness **(C)** Corresponding scores of pictures shown in A. Intestines were graded according to their epithelial thickness as shown in B.

**Figure 2:**
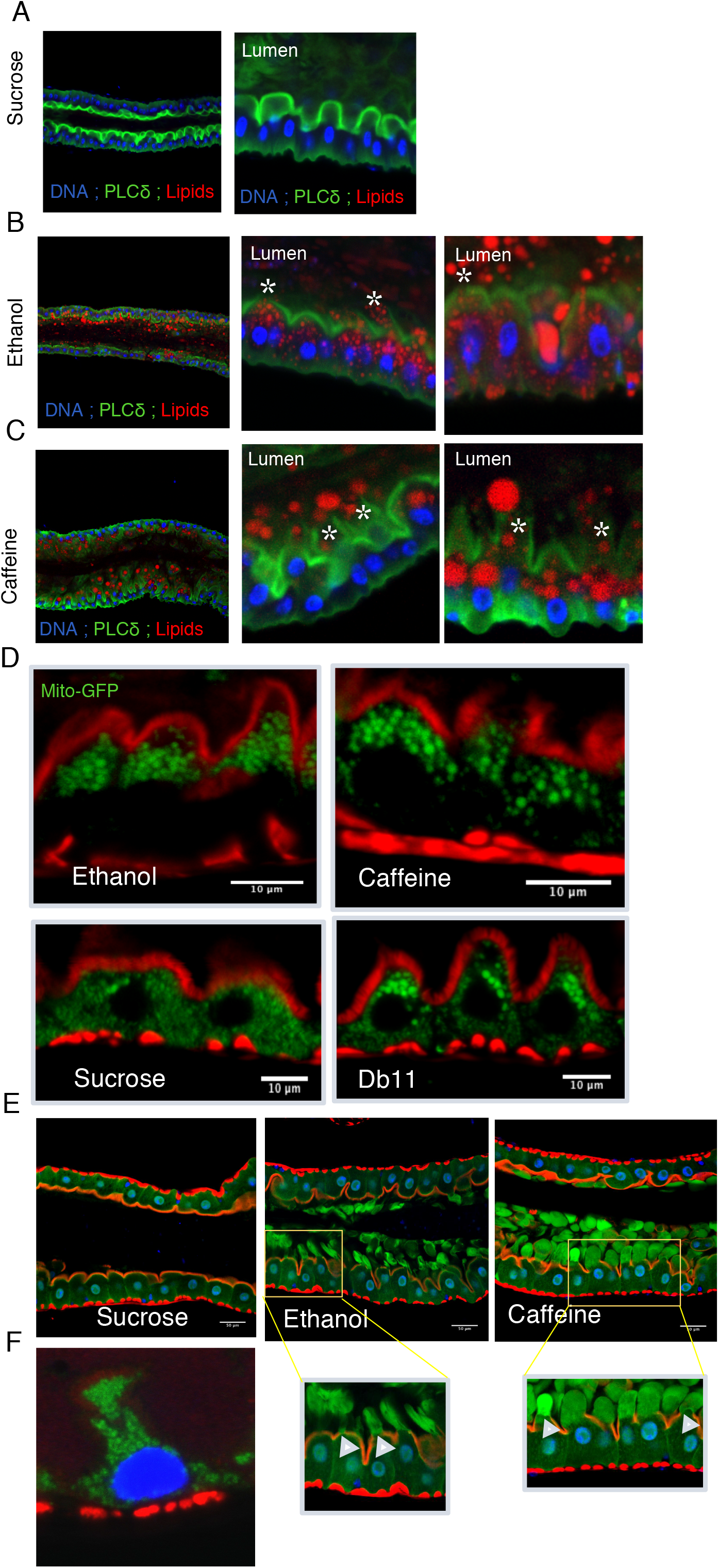
The thinning induced by exposure of enterocytes to caffeine presents the major hallmarks of the apical cytoplasmic purge triggered by *SmDb11*. **(A-C)** Confocal pictures of dissected midguts from NP>PLCδ-GFP flies 1h post-ingestion of 5% ethanol **(B)** or 2,5 mg/mL caffeine **(C)**. Blue = DNA ; Green = PLCδ ; Red = lipids. Compared to sucrose control **(A)**, numerous and large lipid droplets were observed in enterocytes and in the intestinal lumen, upon ethanol **(B)** and caffeine **(C)** ingestion. A disruption of the green (actin) signal was also visible at the site of lipid extrusion (asterisks). *n=30 flies* ; *3 independent experiments*. **(D)** Guts of mito-GFP flies fed either 2.5% ethanol or 2.5mg/mL caffeine were fixed and stained with Texas-Red-Phalloidin. Flies exposed to sucrose solution or *SmDb11* served as negative and positive controls respectively. **(E)** *NP1>UAS-GFP* flies were fed either sucrose, 2.5% ethanol or 2.5 mg/mL caffeine and dissected 30 minutes post-ingestion. Scale bar represents 50μm. A higher magnification of some enterocytes extruding their apical cytoplasm in the boxes is shown below. **(F)** Mito-GFP-labelled mitochondria being extruded from one enterocyte.

### The intestinal epithelium undergoes only one round of thinning during a continuous exposure to SmDb11

Our oral infection paradigm is that of a chronic exposure to *SmDb11* bacteria in sucrose solution on a filter: flies continuously feed on the bacterial solution until the end of the experiment. Here, to ensure that flies were continuously exposed to fresh bacteria, we designed an experiment in which we placed the flies twice a day on a fresh bacterial solution, or sucrose solution for controls, and measured the thickness of the gut epithelia three, seven, and 16 hours after flies being placed on the fresh solution (Fig. 3A). This regimen was established for five days. Afterwards, sucrose solution was added once a day to the filters until an ultimate exposure to fresh bacteria on Day 10. As shown in Fig. 3B-C, we observed a thin epithelium only three and seven hours after the first exposure to *SmDb11*. In contrast, the thickness of the different epithelia of a same batch was varying three hours after exposure to a fresh bacterial solution on the subsequent days, resulting in an increased standard deviation for this time point. The average thickness, however, was not significantly different from controls. Interestingly, the thickness of the epithelia of treated flies 16 hours after exposure to a fresh solution was repeatedly higher every day except on the tenth day than that of controls (Fig. 3D). Importantly, the bacterial gut titer remained constant throughout the experiment (Fig. 3E). Thus, we conclude that under a continuous bacterial exposure paradigm, the gut epithelium undergoes thinning only once, upon the initial exposure.

**Figure 3:**
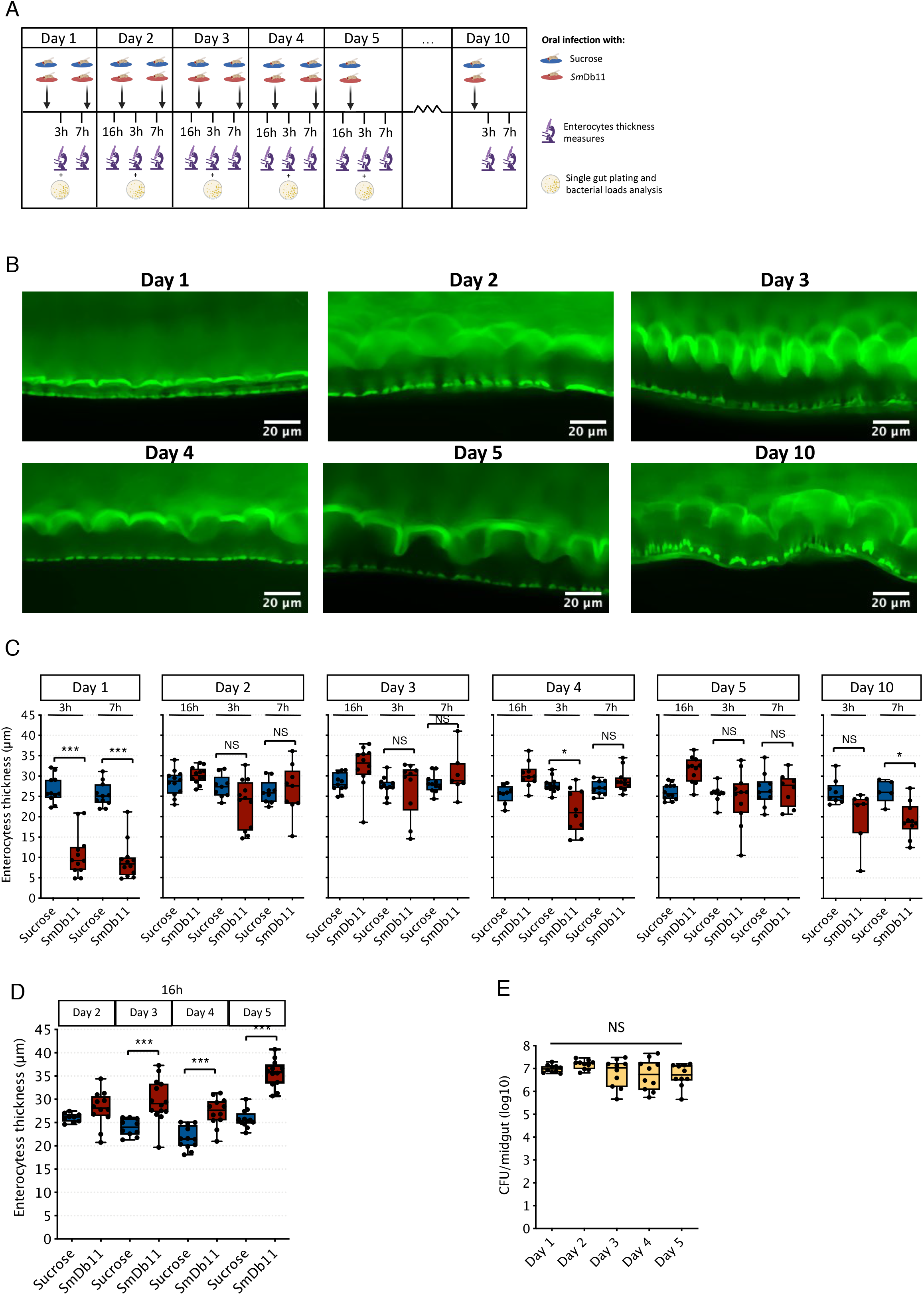
Flies fed chronically *SmDb11* undergo a thinning of the intestinal epithelium only initially upon the first exposure. **(A)** Wild-type flies were either exposed to sucrose or *SmDb11* during 10 days. Fresh bacterial cultures were provided twice a day, in the morning and in the afternoon (except between day 5 to10 during which only sucrose solution was added every day) and intestinal thickness was measured 3h and 7h after the morning exposure and 16h after the evening exposure. The guts were dissected at these time-points to visualize and measure epithelium thickness **(B, C, D)** and at 3h to measure bacterial loads **(E). (B)** Representative confocal pictures of the midguts 3h post infection for each day. Midguts display a very thin epithelium at 3h on Day 1 but not on the following days. **(C)** Enterocyte thickness from the midguts for the time-points described in (A). There is a decrease in epithelium thickness on day 1 at 3h and 7h (*lmer*, \*\*\**p*<0.001), but this decrease is not observed or at least to the same extent in the following days (*lmer*, NS *p*>0.05, \**p*<0.05). **(D)** Thickness of gut enterocytes 16h post infection exposed to a fresh *SmDb11* culture or sucrose on the different days. Enterocytes are bigger 16h post infection compared to the sucrose-exposed flies (*lmer*, \*\*\**p*<0.001). **(E)** Gut bacterial loads are similar 3h post infection with fresh bacterial culture during the different days (*lm*, NS *p*>0.05).

### An initial cytoplasmic extrusion precludes a subsequent one only for a few days

As fresh bacteria are expected to express hemolysin, one would have expected enterocytes to undergo repeated rounds of cytoplasmic extrusion and the epithelium thickness to substantially vary during the infection. As this phenomenon did not happen on a significant scale, we therefore wondered whether an initial round of thinning would induce a state of the intestinal epithelium rendering it impervious to further challenges, a possibility we refer to as priming from the Latin “*primus”*: first. We therefore tested this hypothesis by challenging flies for a limited period of six hours with *SmDb11* or sucrose solution as a control, then placing the flies on regular food until a secondary exposure with fresh bacteria or sucrose solution. As shown in Fig. 4A, the control flies but not the primed flies underwent a thinning of the epithelium. We had already established that this phenomenon also occurs when flies are chronically exposed to *SmDb11*, as shown in Fig. 3 and also in Fig. 4B (*SmDb11* Day1 and *SmDb11* Day 2). This priming phenomenon was also observed in another distinct *Drosophila* wild-type background upon challenge with *SmDb11* (Fig. S2A).

**Fig 4.**
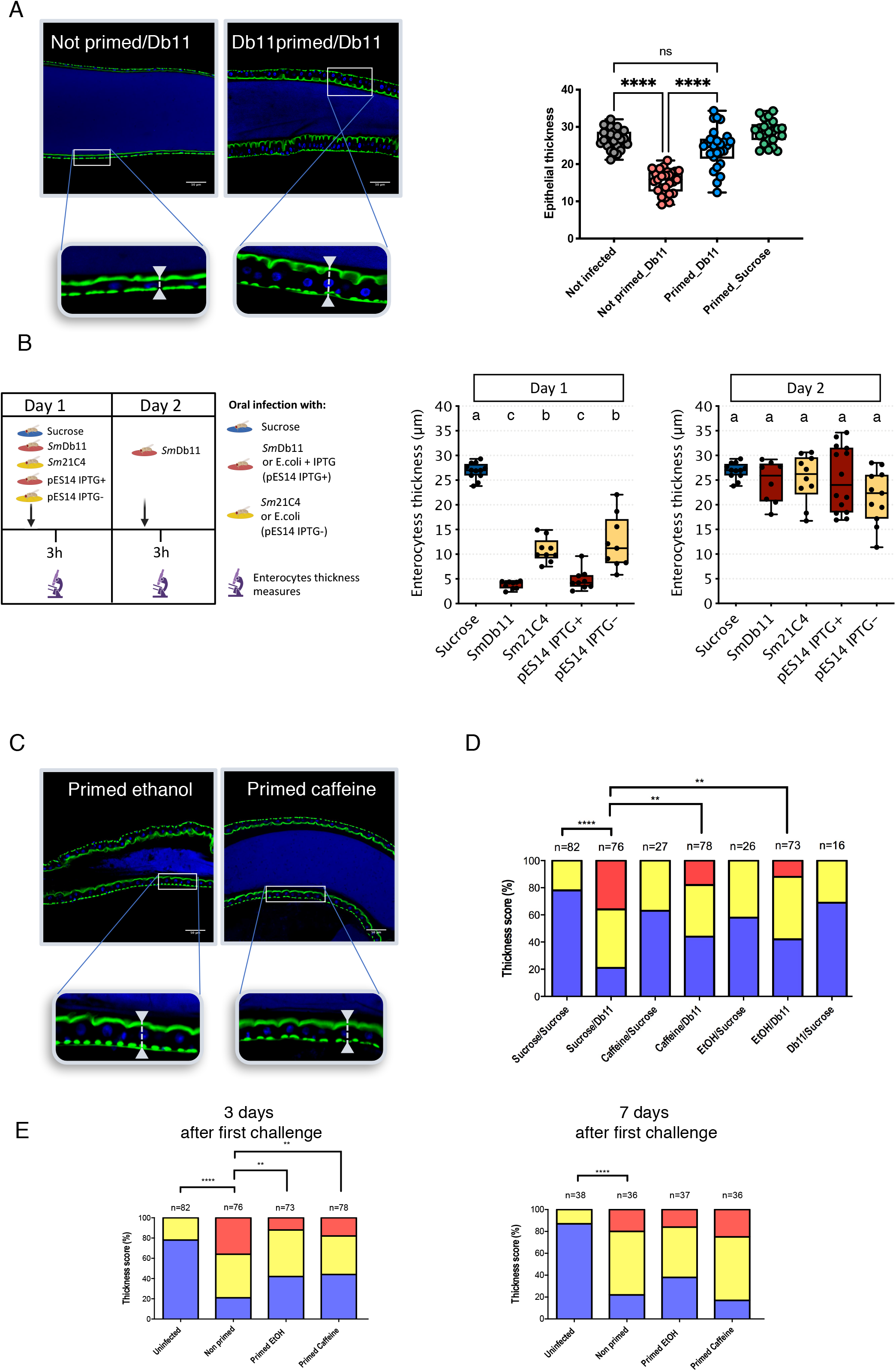
An initial enterocyte cytoplasmic purge triggered by exposure to *SmDb11* or xenobiotics primes the epithelium against thinning upon secondary *SmDb11* challenge Ethanol and caffeine can also prime the enterocytes from an extensive second cytoplasmic purge. **(A-C)** Confocal pictures and zoom of infected R2 midgut region of flies which were exposed to a first 6h-long challenge with sucrose (non-primed control), *SmDb11* (A), ethanol or caffeine (C) prior to being placed again on regular food until they were challenged again close to three days later with *SmDb11*. The quantification for the non-primed controls and *SmDb11* primed flies are shown on the right panel in (A). **(B)** Flies were first exposed to sucrose, *S. marcescens* expressing hemolysin (*SmDb11*), *S. marcescens* mutant for hemolysin (*Sm*21C4), *E. coli* expressing hemolysin (pES14 IPTG+) or control *E*.*coli* (pES14 IPTG-). Flies were exposed to these bacteria on Day 1 and exposed to *SmDb11* at Day 2. Sucrose flies were exposed to sucrose again at Day 2. Epithelium thickness was measured 3h after the first challenge at Day 1 and 3h after the second challenge at Day 2. Hemolysin positive bacteria present a drastic decrease in the gut epithelium thickness at Day1, thgat is distinct from the lesser reduction in the thickness caused by bacteria that do not express hemolysin (*lm*, different letters correspond to statistically significant different groups) (middle panel). All the flies exposed to the four different bacteria on Day 1 become protected from a second cytoplasmic purge after Db11 exposure on Day 2 (right panel). (**D**) Qualitative quantification of the experiment shown in (C) with additional controls. Conditions of the experiment described in (A). Intestines were scored according to their epithelial thickness: thick, semi-thin or thin (Fig. 1B). Statistical tests were performed using χ^2^ test. ****p<0.0001, **p=0.0047 (caffeine), **p=0.0013 (ethanol). Only significant comparisons are shown. (**E**) Right panel: *w*^*A5001*^ flies were exposed to sucrose 100mM, Ethanol 2,5%, Caffeine 2,5mg/mL, or Db11, in order to induce a first cytoplasmic purge. After this first challenge, flies were put back on regular medium. One week after the first exposure, flies were infected with Db11 OD_600_=10 and then dissected 3h after infection. Intestines were scored according to their epithelial thickness: thick, semi-thin or thin. Statistical tests were performed using χ^2^ test. ****p<0.0001, ***p=0.0001, *p=0.0475. Left panel: a subset of quantifications displayed in (D), three days of delay between first and second exposure, is shown again for comparison. Priming lasts for at least two days and is no longer observed by seven days.

We had noticed that older flies behaved more erratically in this assay and therefore compared young seven day-old flies to 21 day-old flies. The older flies underwent a thinning of their epithelia upon a second challenge that was as extensive as after the initial challenge (Fig. S2B). We conclude that older flies have lost the ability to prime their enterocytes against a subsequent challenge.

*SmDb11* mutant flies that are unable to express hemolysin, the 21C4 strain, are able to induce a limited thinning (Lee et al., 2016). We therefore tested whether the exposure to this strain was sufficient to protect the intestinal epithelium from subsequent thinning and found that this lower stimulus was nevertheless sufficient to prime the intestinal epithelium against a secondary *SmDb11* challenge (Fig. 4B, *Sm21C4* and *SmDb11* Day 2). *Serratia* hemolysin produced from *Escherichia coli* harboring the IPTG-inducible pES14 plasmid has been shown to be sufficient to cause epithelial thinning and to be as potent as *SmDb11*(Lee et al., 2016), as illustrated in Fig. 4B (pES14 IPTG+ and *SmDb11* Day1). It was also sufficient to prime the intestinal epithelium against a second exposure to the same stimulus (Fig. 4B, Day 2). Conversely, an initial challenge with *SmDb11* conferred protection against epithelial thinning induced by hemolysin on Day 2 (Fig. S2C, pES14+ Day2). One control for these experiments is to use noninduced *E. coli* bacteria, which unexpectedly were almost as effective as 21C4 to induce epithelial thinning, albeit with more variability (Fig. 4B Day 0). It was also a stimulus sufficient to prime the intestinal epithelium against a secondary exposure to *SmDb11* (Fig. 4B Day 1). One cannot however formally exclude a leaky expression of hemolysin in the noninduced *E. coli*. Fig. S2D establishes that flies challenged by wild-type or hemolysin-deficient *S. marcescens* or *E. coli*, expressing hemolysin or not, recover from the initial bacterial challenge when secondarily exposed to a control sucrose solution.

Interestingly, a primary challenge with caffeine or ethanol was also sufficient to confer a significant degree of protection against a secondary challenge with *SmDb11*(Fig 4C-D). We next determined that the protection conferred by priming with xenobiotics was gained for at least three days but did not last up to seven days (Fig. 4E).

### Identification of genes required for the recovery of initial epithelial thickness in the chronic SmDb11 ingestion model

We have been focusing on the study of the recovery phase by testing multiple genes, for instance those identified by Cronin *et al*. (Cronin et al., 2009) and retaining those that still exhibited a thin epithelium phenotype 16 or 24 hours after a chronic challenge with *SmDb11*. In this way, we have identified the p38b pathway members encoding the Mitogen-Activated Protein (MAP) kinase kinase kinase ASK1, the MAP kinase kinase Licorne and the MAP kinase p38b (Fig. 5A-D). *E5* was a hit of the genome-wide RNAi screen performed on mutant lines feeding on *SmDb11*. It encodes a homeobox-containing transcription factor that is closely related to another such transcription factor encoded by *empty spiracles* (*ems*), as *E5* and *ems* result from a gene duplication event. Interestingly, RNAi lines affecting these genes failed to exhibit a thick epithelium 16 hours after *SmDb11* challenge (Fig. 5E-H).

**Figure 5:**
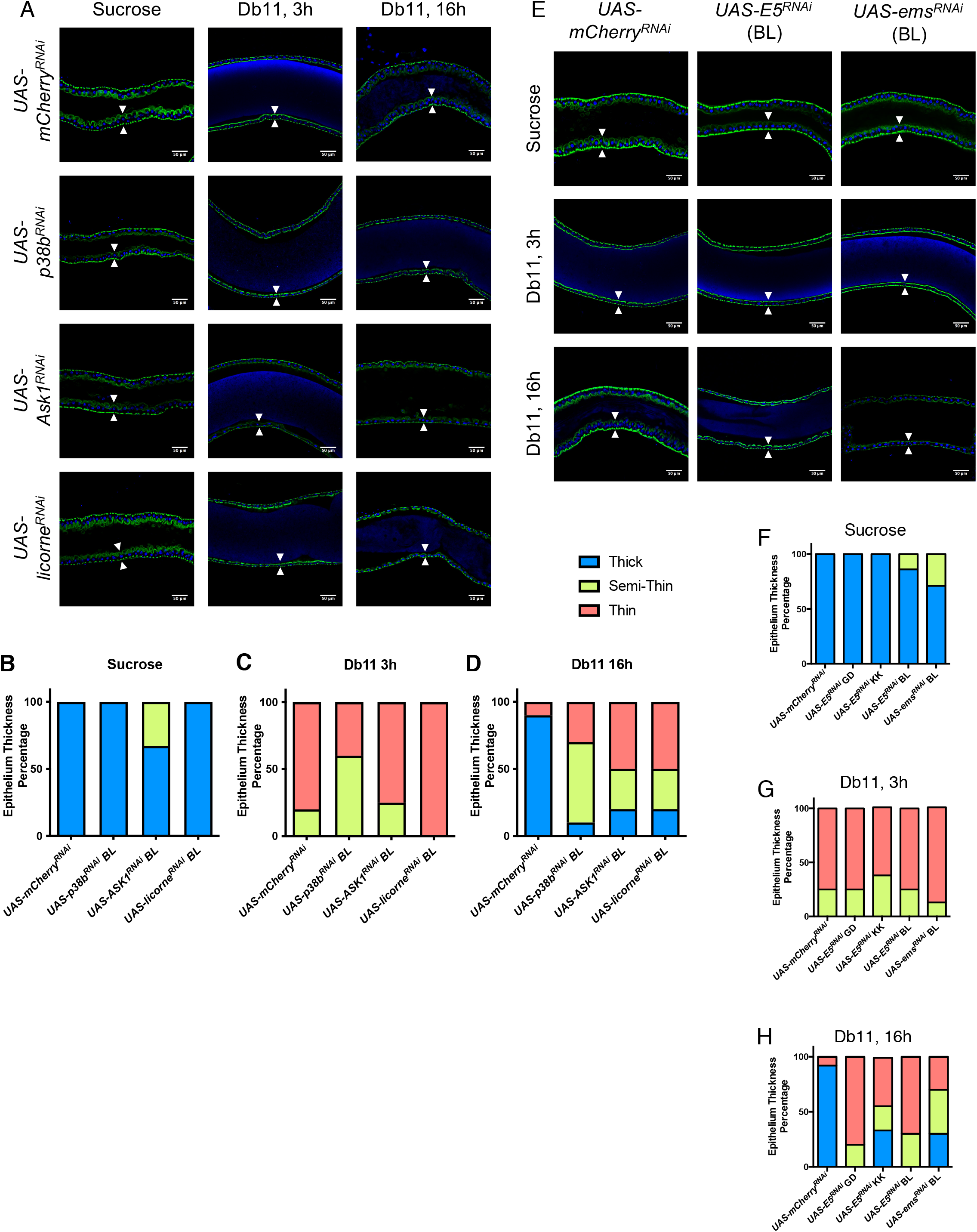
The *p38b* signaling pathway as well as *E5* and *ems* are required for a complete recovery of the intestinal epithelium shape and volume after *SmDb11*-induced thinning. **(A)** Confocal pictures of the R2 region of the anterior midguts of *mCherry*^*RNAi*^, *p38b*^*RNAi*^, *Ask1*^*RNAi*^ and *licorne*^*RNAi*^ mutant flies fed with either sucrose or Db11 (3 and 16h). Actin is stained in green whereas DNA appears blue. **(B-D)** Qualitative quantification of epithelium thickness of midguts (see Fig. 1B) treated with sucrose (B), Db11 3h (C) or Db11 16h (D). **(E)** Confocal pictures of the R2 region of the anterior midguts of *mCherry*^*RNAi*^, *E5*^*RNAi*^ and *ems*^*RNAi*^ mutant flies fed with either sucrose or Db11 (3 and 16h). **(F-H)** Qualitative quantification of epithelium thickness of midguts treated with sucrose (F), Db11 3h (G) or Db11 16h (H). White arrows indicate the thickness of the intestinal epithelium (A and E). Epithelium thickness percentage of flies fed with either sucrose **(A-D)** All crosses *NP1-Gal4Gal80*^*ts*^ x *UAS-RNAi* were performed at 18ºC then the progeny was kept at 29° C (3 days) for the activation of Gal4. Each histogram bar represents values from 10-15 midguts. Each graph represents one out of three independent experiments that yielded similar results.

### The p38b MAPK and E5/ems genes are required for priming and not for the recovery of enterocyte shape

We reasoned that a lack of recovery of the thickness of the intestinal epithelium after cytoplasmic extrusion occurring in enterocytes might result from two distinct events: a lack of recovery of enterocyte shape and volume following the initial cytoplasmic purge or, alternatively, that a process required for priming might be affected and lead to a persistent thinning of the epithelium due to continuous exposure to hemolysin in the chronic ingestion model, thus impairing recovery by a distinct mechanism. To discriminate between these two possibilities, we first performed a series of discontinuous, acute infections in which the flies were exposed only temporarily to *SmDb11* and then placed on sucrose solution. When compared to *p38b, E5* or *ems* RNAi mutant flies continuously exposed to *SmDb11* in the chronic ingestion model, we found that intestinal epithelia of the same genotype exposed only temporarily to the bacteria in the acute infection model recovered a thickness similar to that observed for the *mCherry* RNAi control flies (Fig. 6A). These data suggest that these genes are not required for the recovery of enterocyte shape. We next directly tested the priming itself by exposing control or mutant flies to two *SmDb11* challenges with a period of rest on sucrose solution after the initial three hour-long exposure, which allowed their intestinal epithelia to recover its original thickness (Recovery control, Fig. 6B). As shown in Fig. 6B (Priming panel), flies deficient for *Ask1, p38b, E5* or *ems* were unable to provide substantial protection against the thinning induced by a second *SmDb11* challenge. Thus, the thin epithelium phenotype observed 12 or 24 hours after the beginning of *SmDb11* ingestion is most likely due to impaired priming and not a deficient recovery phase.

**Figure 6:**
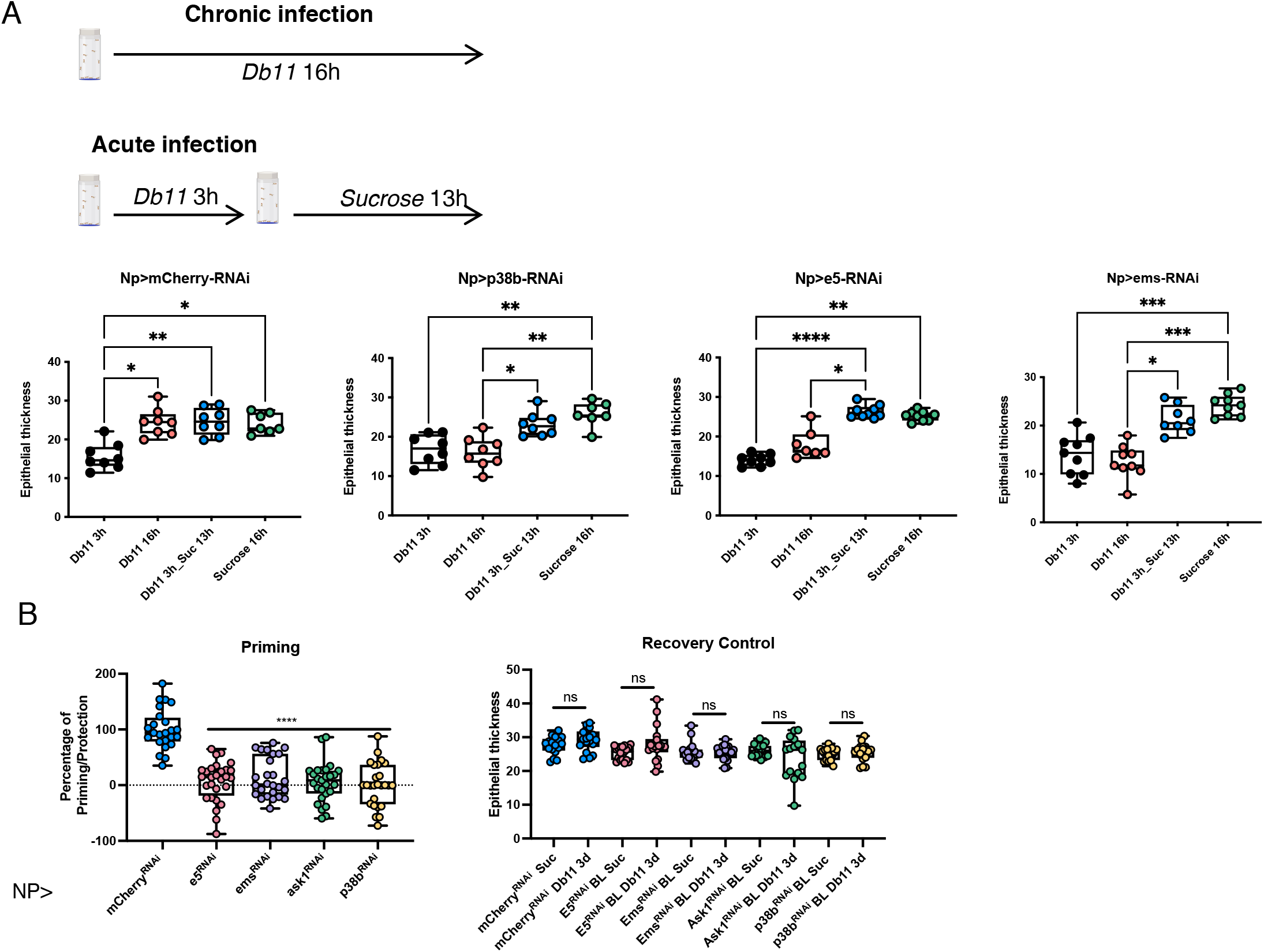
*p38b* signaling pathway and *E5* and *ems* homeobox genes are required for priming against recurring enterocyte cytoplasmic purge and not for the recovery phase that follows the thinning of the *SmDb11*-induced thinning. **(A)** Top panel: experimental scheme: flies are either exposed chronically to *SmDb11* or just for three hours in an acute infection model. The expectation is that a line silenced for a gene required for the recovery of enterocyte shape and volume after cytoplasmic extrusion will present a similar thin phenotype in both types of model. Epithelium thickness measurements of *mCherry*^*RNAi*^, *p38b*^*RNAi*^, *E5*^*RNAi*^ and *ems*^*RNAi*^ mutant midguts that underwent either a chronic infection (Db11 16h) or an acute infection (Db11 3h_Suc 13h). Each dot indicates the average of 10 measures per midgut. Each graph represents one out of three independent experiments that yielded similar results. Statistical analysis was performed using the Kruskal-Wallis test. **(B)** Left panel: Percentage of priming/protection that occurs in the midgut epithelium of the different mutant. The percentage represents the thickness of midguts after a second challenge after normalization with uninfected (sucrose only) and infected (first challenge: *SmDb11* 6h) for each mutant background. This direct assay of priming shows that the tested genes are required to protect the epithelium from recurrent thinning. Right panel: epithelium thickness measurements of mutant midguts at three days comparing the uninfected and infected (first challenge, day 1) and not exposed to a second *SmDb11* challenge. This control experiments checks that the silenced genes are not impinging on long-term recovery of intestinal epithelium thickness. Each graph represents the pooled data from three independent experiments. Statistical tests were performed using One-Way Anova.

**Figure 7:**
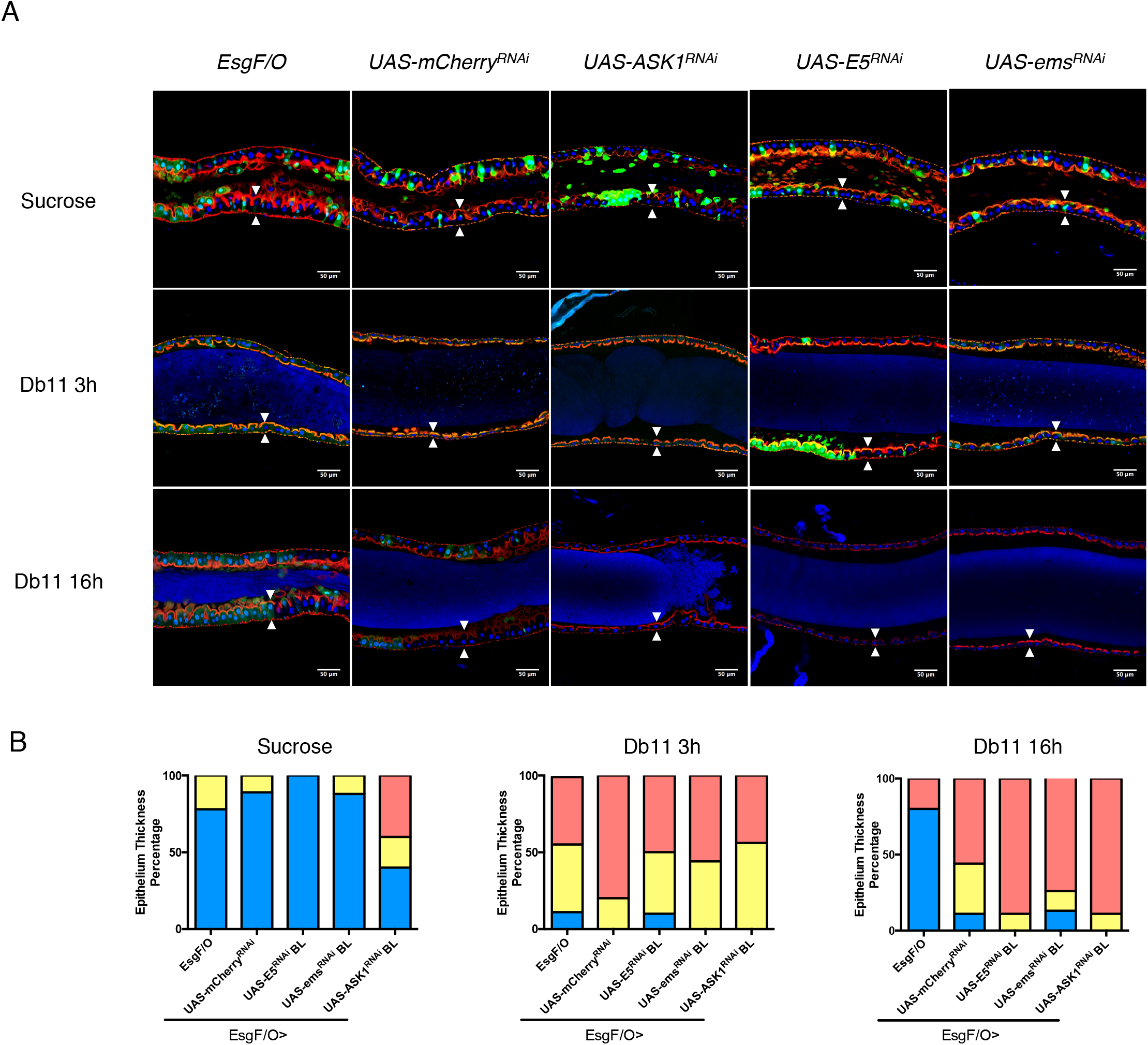
A cell nonautonomous priming role of *Ask1, E5* and *ems*. All crosses *EsgF/O* x *UAS-RNAi* were performed at 18ºC then the progeny was kept at 29° C (3 days) for the activation of Gal4. **(A)** Confocal pictures of R2 region of the anterior midguts of *EsgF/O* (control) *mCherry*^*RNAi*^ (control), *Ask1*^*RNAi*^, *E5*^*RNAi*^ and *ems*^*RNAi*^ mutant flies fed with either sucrose or Db11 (3 and 16h). White arrows indicate the thickness of the intestinal epithelium. **(B)** Qualitative quantification of the epithelium thickness of midguts (see Fig.1B) treated with either sucrose or Db11 (3 and 16h).

We have recently identified the CG1139 amino-acid transporter that is required for the recovery of the original thickness of the intestinal epithelium 16 hours after chronic exposure to *SmDb11* (Socha et al., 2021). When tested in the acute infection assay, the *CG1139* RNAi flies exhibited a recovery that was as limited as that measured after a chronic exposure to *SmDb11* (Fig. S3A). This suggests that *CG1139* is required for the recovery of enterocyte shape and volume even if the epithelium has been exposed to *SmDb11* only for three hours. Next, a priming assay was performed and established that the *CG1139*-deficient flies are protecting the intestinal epithelium against thinning upon a second exposure to the bacteria (Fig. S3B). These data taken together with our study of *CG1139* establish that *CG1139* is a *bona fide* gene required for the recovery phase that follows epithelial thinning and is that thus phenomenologically distinct from the priming process.

We conclude that the p38b MAP kinase pathway and the E5 and EMS transcription factors are required for protection against a secondary exposure to *SmDb11* and that, unlike *CG1139*, they are not required for the initial recovery of the epithelium.

### An unexpected cell-nonautonomous effect of Ask1 and E5/ems following the ingestion of SmDb11

We have previously reported that *CycJ* clones exhibit a cell-nonautonomous effect in that the enterocytes in *CycJ*-deficient clones recover as well as their wild-type neighbors from a continuous exposure to *SmDb11*. We therefore similarly tested the *Ask1, E5*, and *ems* genes using the *esg-Gal4* flip-out technique (Jiang et al., 2009). We noted a much-reduced number of multicellular GFP-labeled clones when *E5* or *ems* RNAi was employed, suggesting a possible role for these genes in the proliferation of gut progenitor cells. Nevertheless, the striking result was that even though the number of wild-type enterocytes was much superior to that of enterocytes present in mutant clones, the whole epithelium comprising wild-type and mutant cells remained thin after 16 hours of chronic exposure to *SmDb11*. We conclude that the *Ask1, E5*, and *ems* genes function in a cell nonautonomous manner to confer protection to the epithelium persistently challenged with *SmDb11*.

## Discussion

The cytoplasmic purge is a fast response to the exposure to microbial pore-forming toxins. Here, we have documented that the ingestion of certain toxicants triggers a similar response of the epithelium. We also demonstrate that a priming phenomenon prevents this fast response from being persistently triggered during a chronic infection thus avoiding the continuous formation of a thin epithelium that is likely not functional. We have identified the apical part of the p38 MAPK pathway and the E5/EMS transcription factors as being specifically required for priming, a cell-nonautonomous process. Thus, the cytoplasmic extrusion of enterocytes appears to be a rapid response to accidental contamination of the ingested food by microbial toxins or xenobiotics.

### A cytoplasmic purge program may be initiated upon exposure to toxins or toxicants

A defining feature of the cytoplasmic purge is the resulting thin intestinal epithelium. This phenomenon may be more rapid than that triggered by exposure to *SmDb11* in the case of ingestion of ethanol or caffeine but may be also delayed in the case of paraquat. Nevertheless, we have observed for these three tested xenobiotics the transient formation of lipid droplets and megamitochondria in the apical cytoplasm, which also occur upon exposure to hemolysin. One possible cause for the fusion of mitochondria into megamitochondria and for the formation of lipid droplets might be oxidative stress, which we have also measured in mitochondria in the case of *SmDb11* infection (Terreri *et al*., in preparation). We also note that we did not observe a significant oxidation after ethanol challenge, which correlates to some extent with an apparent less reproducible formation of megamitochondria after this stimulus.

The response of the organism to ingested xenobiotics is thought to rely on detoxification processes that rely on Phase I&II metabolizing enzymes such as P450’s and Phase III transporters, although the fluidity of membranes is also important in the tolerance to ethanol (Bhaskara et al., 2006; Lieber, 1997; Misra et al., 2011; Montooth, 2006; Rekka et al., 2019; Willoughby et al., 2006). The findings reported here open the possibility that as for PFTs, the cytoplasmic purge may partially protect the organism from the noxious effects of ingested xenobiotics, in addition to classical detoxification pathways.

We have tested a limited number of xenobiotics and at least one of them, rotenone, an inhibitor of complex I of the electron transport chain in mitochondrial respiration, did not appear to trigger a thinning of the epithelium. In the case of paraquat and Cd, we have not demonstrated that there is indeed an extrusion of the apical cytoplasm even though we did detect the formation of megamitochondria that appeared early in the case of paraquat exposure even though a thin epithelium was only observed hours later.

Taken together, these data suggest the existence of a common process that is triggered by exposure to these stimuli. Indeed, given the large apertures that form to expel the apical cytoplasm, it appeared unlikely that this would be caused by *SmDb11* hemolysin that forms only pores of 2-3 nm diameter (Hertle, 2000, 2005). In mammalian macrophages, the activation of the inflammasome has been reported to lead to the formation of openings in the cytoplasmic membrane through the activation of the PFTs gasdermins and also the involvement of ninjurin1 that is required for the formation of large apertures that initiate pyroptotic cell death (Churchill et al., 2021; Kayagaki et al., 2021; Ruhl and Broz, 2021). Whereas gasdermin homologs have not been identified in the *Drosophila* genome, there are three *Drosophila* Ninjurins. We have tested so far only *NinjurinC* by RNAi knock-down, without success. However, we have identified at least one caspase required for the initiation of the cytoplasmic purge. Ongoing studies will allow to determine whether the evolutionarily conserved cytoplasmic purge mechanism rely on a conserved mechanism akin to that of the inflammasome.

### Priming, a safeguard to prevent the chronic activation of the cytoplasmic purge defense mechanism

We had observed early on that the overall structure of the midgut epithelium appeared to be relatively well-preserved during chronic *SmDb11* infections (Nehme et al., 2007). One possibility would have been that bacteria placed on the filter in sucrose solution might no longer produce hemolysin. Hence, we have excluded this possibility in this study by providing fresh bacteria every day. Indeed, when priming is impaired the epithelium remains thin, which has also been observed in *CycJ* mutants that rapidly succumb to the ingestion of *SmDb11* (Lee et al., 2016). It is likely that a thin epithelium is dysfunctional and would be unable to sustain digestion over the long term even though biochemical assays revealed only a slow loss of digestive enzyme activity in starving flies. The existence of priming implies that the cytoplasmic purge program has likely been selected to deal with accidental contaminations of the ingested food and may be counter-productive in the case of chronic infections.

The study of the thin epithelium phenotype induced by the ingestion of *SmDb11* has been hampered by a decreased or altogether absent phenotype that has been occurring for more or less prolonged periods over the years throughout our studies of this phenomenon. For one line that appeared to be resistant to thinning, we finally determined that a strain of *S. marcescens* was a constituent of is gut microbiota and might account for the resistance since this line became permissive again after bleaching the eggs to remove the microbiota. It may have been that this line was primed against any further experimental challenges. In contrast, we suspect a contamination of our food or water supply when this phenomenon was affecting all of our experiments, even though we were never able to identify the origin of the problem. One exception is when our laboratory tested another recipe of fly food and used an alcoholic formulation of the food preservative whereas our traditional formula relied on the use of salts of the same preservative: this allowed us to identify ethanol as being a chemical able to induce intestinal thinning as well as priming.

The exact mechanism of priming remains elusive at this stage and will likely require a fine-grained understanding of the purge process to identify which step is affected by priming. At the enterocyte level, we observed a correlation between the limited duration of the priming and the life expectancy of enterocytes in *SmDb11*-infected guts (the compensatory proliferation of ISCs takes place 24 hours after challenge in the case of chronic infections as it is no longer prevented by the cytoplasmic purge that limits the damages inflicted to enterocytes (Lee et al., 2016)). However, the involvement of some members of the p38b stress-response pathway that appears to function in a cell-nonautonomous manner indicates that priming is likely to be a complex process.

### Cell-autonomous and cell-nonautonomous processes involved in priming

Our genetic analysis identified the distal part of the p38 MAPK pathway on the one hand and the transcription factors E5 and EMS on the other to be required for the priming phenomenon. Indeed, whereas mutant RNAi lines targeting these genes yielded the same phenotype when comparing chronic *vs*. acute infection, that is a recovery only when flies were placed on food after a limited exposure to *SmDb11* (Fig. 6A), they failed to provide any protection against a second exposure to *SmDb11* in the priming assay (Fig. 6B). The ASK1-Licorne-p38b axis is likely to function in a linear chain as described for the response to other stresses. P38b is thought to activate transcription factors of the ATF family. We however failed to reveal any reliable “recovery” phenotype when we tested *ATF2, ATF3, ATF4* or *ATF6* by RNA interference. This also applied to *HR38*, a nuclear family receptor homologous to mammalian NR24A2 (NUR77) (Watanabe et al., 2015). ASK1 can be activated by oxidative stress and might therefore provide a link to connect the mechanism triggering the purge and that involved to mediate the priming effect. ASK1 has been described to be positively or negatively modulated by many interactors (Takeda et al., 2008), some of which we have tested. Interfering by RNAi with the expression of Trx-2, 14-3-3ζ or ε, TRAF-like, a TRAF2 fly homolog, HR38 or PpD3 failed to reveal any phenotype after exposure to *SmDb11*, either at three hours or 16 hours. Indeed, we expected to reveal in the case of negative regulators a constitutive inhibition of the cytoplasmic purge. This may indicate that the priming phenomenon may also involve a second pathway that also needs to be activated in conjunction with the p38b MAPK pathway.

Interestingly, we have identified two genes encoding homeobox transcription factors, *E5* and *ems*, that display properties similar to those of genes of the p38b signaling pathway. Homeobox-containing transcription factors may heterodimerize through this domain (Burglin and Affolter, 2016; Zaffran and Frasch, 2005). This property might account for the shared phenotypes of *E5* and *ems*, which was not enhanced by knocking down both genes simultaneously.

We have been unable to determine whether the p38b MAPK pathway genes and the *E5/ems* genes function in the same pathway for lack of good tools for epistasis analysis. As related above, affecting the *Trx-2* or *PpD3* genes that code for negative regulators of ASK1 did not reveal a dominant phenotype. The overexpression of p38b pathway genes was also unfruitful. We note however the presence of a consensus p38b phosphorylation site on CyclinJ, which itself might be able to interact with E5, at least in a bacterial two-hybrid system (Stanyon et al., 2004). Thus, CycJ may provide a link between the p38 MAPK pathway and E5/EMS. Interestingly, all of these genes display a thin epithelium phenotype 16 hours after chronic exposure to *SmDb11*. Interestingly, we have observed that this thin midgut epithelium phenotype disappears upon acute infection and subsequent resting period on regular food. Unfortunately, *CycJ* knocked-down flies have displayed so far a phenotype that is too variable in priming assays to make a reliable conclusion on its involvement in the priming process (ST, unpublished data). Thus, further work is required to clarify this important point.

In any case, it is striking that clonal analysis performed with either p38MAPK pathway genes and *E5/ems* genes on the one hand and *CycJ* on the other yield both non-autonomous phenotypes, but with opposite outcomes. Thus, should CycJ be a link between p38b and E5/EMS, it might be inhibited by p38b-mediated phosphorylation and thereby relieve a potential suppression of E5/EMS transcription activation function. The finding of an epithelium remaining thin when only a limited number of cells is mutated for either the *p38b* pathway or *E5/ems* is puzzling. It suggests that these genes are required to prevent the expression of a diffusible inhibitor of priming that would be released upon the ingestion of *SmDb11*. The identification of such a molecule will constitute a major challenge to an in-depth understanding of the priming phenomenon that protects the intestinal epithelium from unwanted side-effects of a host defense likely developed to deal with occasional encounters with microbial toxins or toxicants.

## Material and Methods

### Fly husbandry and fly stocks

*Drosophila melanogaster* flies were raised at 25°C and nearly 60% humidity, 14h of daylight and fed a standard semi-solid cornmeal medium (6.4% (w/v) cornmeal (Moulin des Moines, France), 4.8% (w/v) granulated sugar (Erstein, France), 1.2% (w/v) yeast brewer’s dry powder (VWR, Belgium), 0.5% (w/v) agar (Sobigel, France), 0.004% (w/v) 4-hydroxybenzoate sodium salt (Merck, Germany)).

All experiments were performed on three to seven days old female flies, unless specified otherwise.

The wild-type flies used for experiments were white *wA5001* (Thibault et al., 2004) and the DrosDel *w*^*1118*^ isogenic stock (*w*^*1118*^ *iso*) (Ryder et al., 2004); the *GD* line (VDRC stock #60000) and *KK* line (VDRC stock #60100) were used as controls for some RNAi experiments. The following tissue- or cell-type-specific driver lines were used: enterocyte-specific driver line NP1, (also known as Myo31D), *NP-Gal4-tubGal80*^*ts*^ (Cronin et al., 2009; Nehme et al., 2007), progenitor stem cells-specific *esg-Gal4Gal80*^*ts*^, and the flip-out line *w; esgGal4tubGal80*^*ts*^ *UAS-GFP;UAS-flp Act>CD2>Gal4* (Jiang et al., 2009) for the clonal analysis. The roGFP2 flies strains were a kind gift of Jörg Großhans (Albrecht et al., 2011); cytoplasmic-Orp1 (P015, pCasPeR4-cyto-roGFP2-Orp1 II HV (line 1); mitochondrial-Orp1 (P018, pCasPeR4-mito-roGFP2-Orp1 II HV (line 6).The RNAi lines used were from the Vienna Drosophila Research Center (VDRC) or were Bloomington Trip lines (BL): *UAS-mCherry*^*RNAi*^ (BL #35785) *UAS-E5*^*RNAi*^ (VDRC *GD* #47791) (VDRC *KK* #110804) (BL HMJ21631), *UAS-ems*^*RNAi*^ (BL #28726), *UAS-p38b*^*RNAi*^ (BL #35252), *UAS-ASK1*^*RNAi*^ (BL #32464) and *UAS-licorne*^*RNAi*^ (BL #31643). The fluorescent membrane-bound GFP reporter line was: *y*^*1*^*w*^*^; *L*^1^/CyO; P{UAS-PLCδ-PH-eGFP}3/TM6B,Tb^1^ (BL #39693).The UAS-mito-GFP line (import mitochondrial sequence of human cCoxVIII fused to GFP) was a kind gift from Thomas Rival (Rival et al., 2011).

### Microbiology and Infections/exposure to xenobiotics

*Serratia marcescens* strain Db11 (*SmDb11*) was cultured on Lysogeny Broth (LB) agar plates with 100 μg/mL of streptomycin. The strain 21C4 *Sm*21C4 (Kurz et al., 2003) was cultured on LB-agar plates with 20 μg/mL chloramphenicol. The solid plates were placed overnight at 37°C to obtain colonies. For liquid cultures, one bacterial colony was taken off the solid plate and inoculated into 200 mL of liquid LB. These cultures were kept overnight at 37°C with agitation. All infections were performed using a final bacterial OD 600 (Optical Density at 600 nm) of 10, except for the survival where OD=1 was used. The OD was measured with a spectrophotometer and 50 mL of this solution was centrifuged at 4000 g/rcf for 10 minutes. The pellet was re-suspended in the appropriate volume of a 50 mM sucrose solution containing 10% of LB, in order to reach a final OD=10. 2 mL of this infection solution were added to two absorbent pads (Millipore AP1003700) that were placed at the bottom of medium-size vials (3.5 cm diameter). Twenty female flies of five to seven-day old were fed this infection solution, or sucrose 50 mM as a control at 29°C. 2,5mg/mL caffeine solution was prepared by diluting caffeine (Sigma-Aldrich, C0750) in 50mM sucrose and placing the solution for 30 minutes in a 50°C water bath. Paraquat (Sigma-Aldrich 856177) and Cd solutions were prepared in a similar manner by diluting the powder in 50mM sucrose. EtOH final concentrations were either 2,5% or 5% in 50 mM sucrose. 50 mM sucrose was used for control. Survival assay in biological triplicates of 20 flies per vial were performed on three to seven days old flies, at 29°C with 70% humidity,. Survivors were counted each day, and 200μL of sucrose 100mM was added to all the vials, unless otherwisde specified.

### Histology and immunofluorescence

Midguts were dissected in PBS and fixed for 30 minutes with 4% paraformaldehyde. Samples were washed three times with PBS-Triton X-100 0.1% (PBT 0.1%).

For actin staining, midguts were incubated for 1h30 at room-temperature or overnight at 4°C in 10 μM Fluorescein Isothiocyanate (FITC) (Sigma-Aldrich #P5282) or Texas-Red labeled phalloidin (Invitrogen TM #T7471). Samples were then washed three times with PBT 0.1%.

#### Phosphohistone3 (PHH3) staining

The anti-PHH3 antibody (Millipore ref 09-797) was diluted 1:500 in PBT 0,1% + BSA 2%. Samples were incubated in this solution 2h at RT or overnight at 4°C, then washed three times in PBT 0,1% and incubated with goat anti-rabbit FITC antibody (Abcam #6717) diluted 1:1000 in PBT 0,1% + BSA 2%. Samples were finally washed three times in PBT 0,1% prior to mounting.

#### Mounting

All samples were mounted on 8-wells diagnostic microscopy slides (Thermo Fisher Scientific) with Vectashield containing DAPI (Vector Laboratories) and then stored in the dark at 4°C.

#### Microscopy

Samples were observed using a LSM780 confocal microscope (Zeiss) or an epifluorescence Axioscope2 microscope (Zeiss) as needed. Images were taken using a plan/apochromat 20x/0,8 dry objective. Raw files were treated and analyzed using the ImageJ/Fiji software if needed.

### Classification of gut epithelium thickness

To determine the level of thinning and the recovery capacity, we used either qualitative or quantitative analysis. For qualitative analysis, midguts were classified into three different categories according to their epithelial thickness. Thick epithelium, which we represent in blue, corresponds to cells with the normal size, (around 20μm in length), with a clear dome shape characteristic of the intestine. Thin epithelium, which we represent in red, corresponds to cells that are very thin (around 5μm) and the normal dome shape of the cells is not observed. Semi-thin epithelium, which we represent in yellow, corresponds to cells with intermediate thickness (around 13μm), where the dome shape is still not completely defined. For quantitative analysis, pictures of midguts were acquired using Fluorescence Axioskope Zeiss and the thickness of gut enterocytes was measured using ImageJ/Fiji software. The measure for each midgut corresponds to the average of the measure between 10 enterocytes. Measures were taken every 5 to 10 enterocytes.

### Priming experiments

Flies were first exposed to either sucrose or Db11 (OD_600_ 10) for 6h, then shifted to regular fly food. In the third day, flies underwent a second challenge with Db11 or a treatment with sucrose for 3h. In some experiments as noted, the flies were exposed to the bacterial solution for 24 hours and then exposed to a second challenge. The midguts were dissected, fixed with PFA (8%) and stained with phalloidin. The samples were then observed and imaged using the Axioscope 2 microscope (Zeiss). The epithelium thickness of each midgut was measured using FIJI software and the average of 10 measurements per midgut was plotted.

The index of protection is introduced to quantify the extent of protection conferred by a first challenge to a fly line. The principle is to measure the possible extent of thinning of a line by measuring the average epithelial width of naive flies exposed to the second *SmDb11* challenge and to subtract from this average “naive” value the average epithelial thickness of flies of the same line that have been exposed to both an initial and a second *SmDb11* challenge. This difference should thus be low for mutants unable to prime since the epithelium remains thin after the second challenge in this case. Finally, the data are normalized by the values obtained by following the same procedure with reference control flies *NP1-Gal4-Gal80*^*ts*^*>UAS-mCherry*^*RNAi*^.

### Ratiometric roGFP2 probe measurements

The original detailed protocol is found in (Barata and Dick, 2013). We provide below a shortened version.

#### Reference samples

To obtain fully oxidized and fully reduced samples, midguts were dissected in PBS 1x and then incubated 10 minutes in diamide (DA) or dithiothreitol (DTT) for respectively fully oxidized and reduced references. After washing in PBS, these samples were placed 15 minutes in N-ethyl-maleimide (NEM) 20mM in order to preserve the redox state of the tissue. Midguts were quickly washed again in PBS and then incubated in PFA 4% for 20 minutes and mounted in Vectashield without DAPI.

#### Experimental samples

For dissection of experimental conditions samples, PBS was supplemented with NEM 20mM to preserve the redox state of the tissue. The midguts were further alkylated in NEM for 15 minutes after dissection. After washing in PBS, samples were fixed in PFA 4% for 20 minutes and mounted in Vectashield without DAPI. Prior to image acquisition, laser intensity for 405nm and 488nm excitation was adjusted using the fully oxidized and fully reduced samples, respectively. These settings are no longer modified for all the samples in the same experiment. For imaging, the probe was excited sequentially at 405nm and 488nm and emission was recorded at 500-530 nm.

#### Image processing

Image treatment was performed adapting the protocol from (Kardash et al., 2011). Briefly, rolling ball procedure set to 50 pixels was used to subtract background and images were converted to 32-bit. The image intensity was set up with the default setting thresholds in greyscale and dark background. Values below the threshold were set to “not a number”. The plugin RatioPlus was used to generate ratio images by dividing the 405nm picture by the 488nm corresponding picture. The lookup table “fire” was used for final coloring. The intensities were finally normalized to the DTT control (fully reduced).

### Statistical analysis

Statistical analyses were performed using GraphPad software Prism 6 and R. For qualitative analysis of epithelium thickness, we used chi-square statistical tests. We also used Linear models (lm), or linear mixed-effect models (lmer package lme4) if there were random factors (Bates et al., 2015). Significance of interactions between factors was tested by comparing models fitting the data with and without the interactions using analysis of variance (anova). Models were simplified when interactions were not significant. Pairwise comparisons of the estimates from fitted models were analyzed using lmerTest, lsmean, and multcomp packages. The Logrank test was used for survival statistics.

## Acknowledgements

We thank the Vienna Drosophila Resource Center (VDRC, www.vdrc.at) and Bloomington Drosophila Stock Center (NIH P40OD018537) for their resource, Jörg Großhans and Thomas Rival for fly stocks.

We thank Tanja Seissler for performing some early experiments.

This work has been funded by CNRS, University of Strasbourg, ANR grant ANR-16-CE13-0011-01 (ENTEROCYTE_PURGE_RECOVERY), Fondation pour la Recherche Médicale (Equipe FRM DEQ20090515394 to DF) and fellowships FDT201904008233 to ST & FDT20170437224 to CS. Our work is also partially sponsored by Infinitus, Inc (China). The funding sources had no role in the design of the study nor in its execution, analyses, interpretation of the data or decision to publish the results.

