## Supplementary Figures for "A prior exposure to *Serratia marcescens* or xenobiotics primes *Drosophila* enterocytes against a recurring cytoplasmic purge"

**Fig. S1: Phenotypes observed after the ingestion of xenobiotics**

**(A)** Mitochondrial-Orp1-roGFP2 (left panel) and cytoplasmic-Orp1-roGFP2 (right panels) reporter fly lines were treated with sucrose 100mM, EtOH 2,5% or caffeine 2,5 mg/mL. Midguts were dissected 30min after the ingestion. Each dot on the graph represents one intestine. Midguts were also exposed to DTT 20mM or DA 2mM to estimate their responsiveness to reduction and oxidation, respectively. The graph represents the pooled results of three independent experiments. Columns and error bars represent the mean and the SD. For pair-wise comparison, statistical analysis was performed using unpaired t test with Welch's correction (left panel), or Mann-Whitney test (right panel), comparing each condition to sucrose treatment. ****$p<0.0001$. Only significant comparisons are shown.

**(B-C)** Lipid droplets are extruded in the intestinal lumen upon exposure to ethanol or caffeine. Confocal stack of dissected midguts from NP>PLCδ-GFP flies 1h post-ingestion of 5% ethanol (B) or 2.5 mg/mL caffeine (B). Blue = DNA ; Green = PLCδ ; Red = lipids.

To build a virtual endoscopy, samples were imaged with a confocal microscope in 3D by generating a stack of 50 horizontal optical sections separated by 0,5 μm and reconstructed in an orthogonal view with Fiji (ImageJ).

**(D-E)** Survival curves of $w^{A5001}$ flies exposed to xenobiotics and to Db11. (E) Same as (B), but in this experiment ethanol was added each day and bacterial load was decreased to $OD_{600}=1$. Error bars represents the standard error. Statistics were performed using Logrank.

**(F)** $PHH3^{+}$ cell count in midguts of $w^{A5001}$ flies after 24h of exposure. The graph represents pooled results of three independent experiments. Each point represents an individual midgut. Note that exposure to the insecticidal caffeine does not induce the compensatory proliferation of ISCs.

**(G-H)** Confocal micrographs of actin-stained (red, except third row of G, green) mito-GFP (green) flies or wild-type flies (third row of G, lipid- droplets stained red) during 100 mM Cd treatment (G) or 5mM paraquat treatment (H). Note thinning at a late time point (7h) despite early megamitochondria formation. Top panels show a low magnification (scale bars: 50µm) of part of the R2 region; the bottom panels display a higher magnification of enterocytes (scale bars: 10µm).

Error bars show mean with standard deviation (SD). Statistics were done using Kruskal-Wallis test for multiple comparisons, comparing each condition to sucrose treatment. *p<0,05, ***p<0,001, ****p<0,0001. Only significant comparisons are shown.

**Fig S2. Priming of gut enterocytes in another genetic background and absence of priming in older flies**

**(A)** $w^{1118}$ isogenic flies were exposed on Day 1 and Day 2 to fresh Db11 cultures. There is a decrease in epithelium thickness at Day 1 but not at Day 2 (lm, NS p>0.05, *** p<0.001).

**(B)** Seven and 21 days old flies (Young flies and Old flies, respectively) were exposed to fresh solutions of either *SmDb11* or sucrose at Day 0 and Day 1 to access the enterocyte thickness at 3h, 7h and 16h post infection (left panel). Both young and older flies exhibit a decreased thickness of the gut epithelium at 3h and 7h on Day 1 (lm, *** p<0.001). On Day 2, young flies have recovered a normal gut epithelium thickness (lm, NS p>0.05), but this decrease is still observed in older flies, which are not primed against a second bacterial exposure (lm, *** p<0.001).

**(C)** Flies were exposed to *SmDb11* on Day 1 and exposed to *SmDb11* or *E. coli*-expressing hemolysin on Day 2. Both bacteria cause a decrease in the epithelium thickness on Day 0 (*SmDb11* and pES IPTG+) compared to sucrose control. The flies exposed to a first *SmDb11* challenge on Day 0 are primed against a second challenge on Day 2 with either *SmDb11* (SmDb11- *Sm*Db11) or hemolysin-expressing *E. coli* (SmDb11 - pES IPTG+) (*lm*, different letters correspond to statistically significant different groups). These data are the same as those shown in Fig. 4C, except that *Sm*Db11 - pES IPTG+ data are included in this panel. It was plotted together for comparison purposes.

**(D)** Flies were first exposed to sucrose, *S. marcescens* expressing hemolysin (*Sm*Db11), *S. marcescens* mutant for hemolysin (*Sm*21C4), *E. coli*-expressing hemolysin (pES14 IPTG+) and *E. coli* (pES14 IPTG-). Flies were exposed to these bacteria on Day 1 and exposed to Sucrose on Day 2. This is a control for Fig. 4B. Flies exposed to Db11 on the first day present a significantly thicker epithelium 3h after being exposed to sucrose in the second day (*lm*, different letters correspond to statistically significant different groups).

**Fig S3. The amino acid transporter CG1139 is required for recovery and not for enterocyte priming.**

**(A)** *iso*> *CG1139* RNAi or *NPiso*> *CG1139* RNAi flies were exposed to either a chronic infection with *SmDb11* for 16h or to an acute infection for 3h and then changed to sucrose. Epithelium thickness was measured 3h and 16h post infection for both conditions. Control flies recover in both acute and chronic exposure to *SmDb11* while *NPiso*> *CG1139* RNAi present a thinner epithelium at 16h compared to sucrose fed flies for both conditions (*lmer*, NS $p>0.05$, *** $p<0.001$) **(B)** *NPiso*> *CG1139* were tested in a priming assay. Flies were subjected to a first challenge with *SmDb11* (Primed Db11) or to sucrose control (Not primed) for 6h. After this period, flies were transferred to vials containing standard food and were exposed to a second *SmDb11* challenge on the third day. *NPiso*> *CG1139* RNAi flies previously exposed to *SmDb11* have similar epithelium thickness than sucrose-exposed flies and do not present the same decrease observed in non-primed flies (*lm*, NS $p>0.05$, *** $p<0.001$). Thus, *CG1139* is not required for priming.

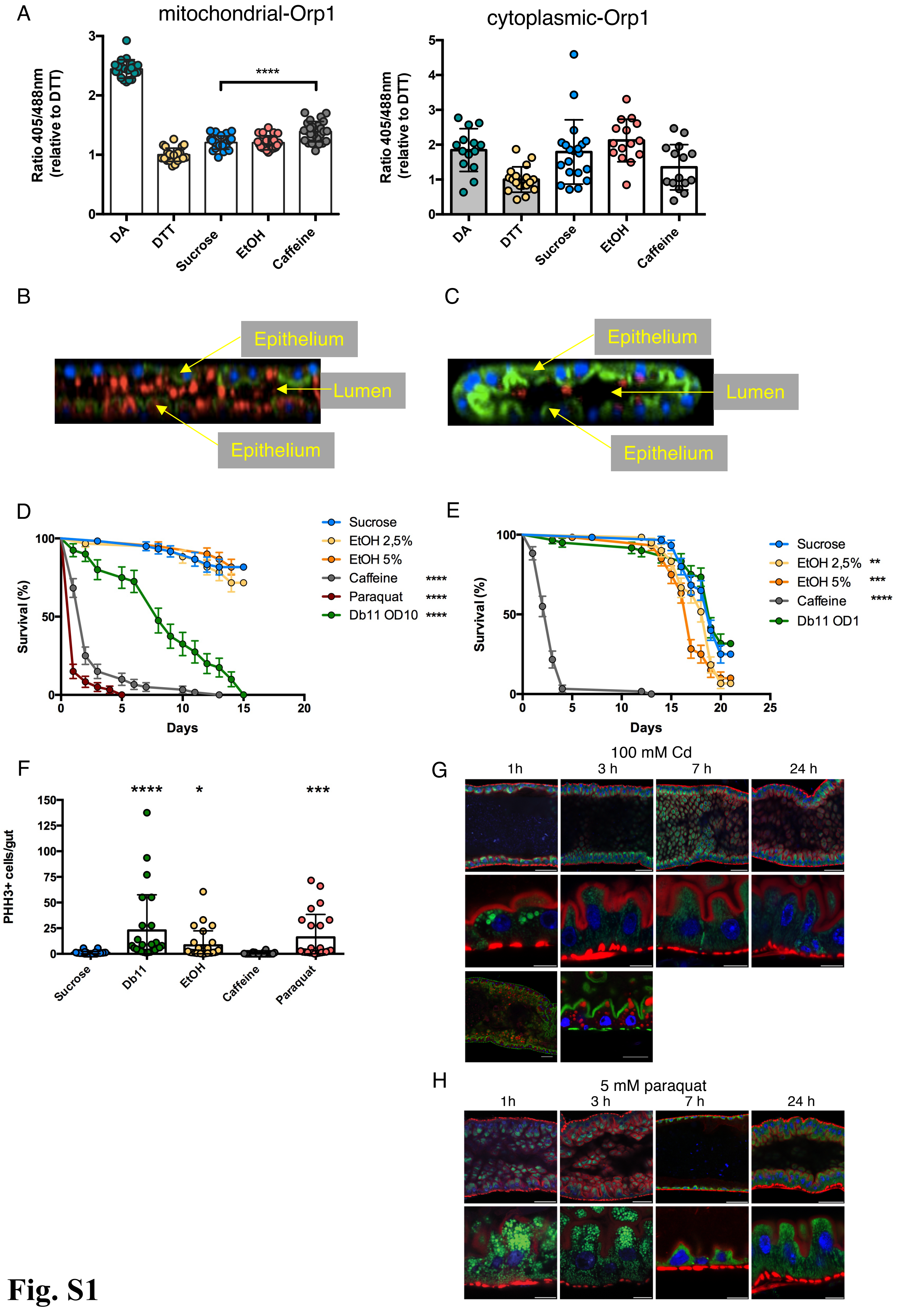
A
mitochondrial-Orp1
Ratio 405/488nm (relative to DTT)
****
DA
DTT
Sucrose
EtOH
Caffeine
cytoplasmic-Orp1
Ratio 405/488nm (relative to DTT)
DA
DTT
Sucrose
EtOH
Caffeine
B
Epithelium
Lumen
Epithelium
C
Epithelium
Lumen
Epithelium
D
Survival (%)
Days
Sucrose
EtOH 2,5%
EtOH 5%
Caffeine ****
Paraquat ****
Db11 OD10 ****
E
Survival (%)
Days
Sucrose
EtOH 2,5% **
EtOH 5% ***
Caffeine ****
Db11 OD1
F
PHH3+ cells/gut
**** * ***
Sucrose
Db11
EtOH
Caffeine
Paraquat
G
100 mM Cd
1h
3 h
7 h
24 h
H
5 mM paraquat
1h
3 h
7 h
24 h

**Fig. S1**

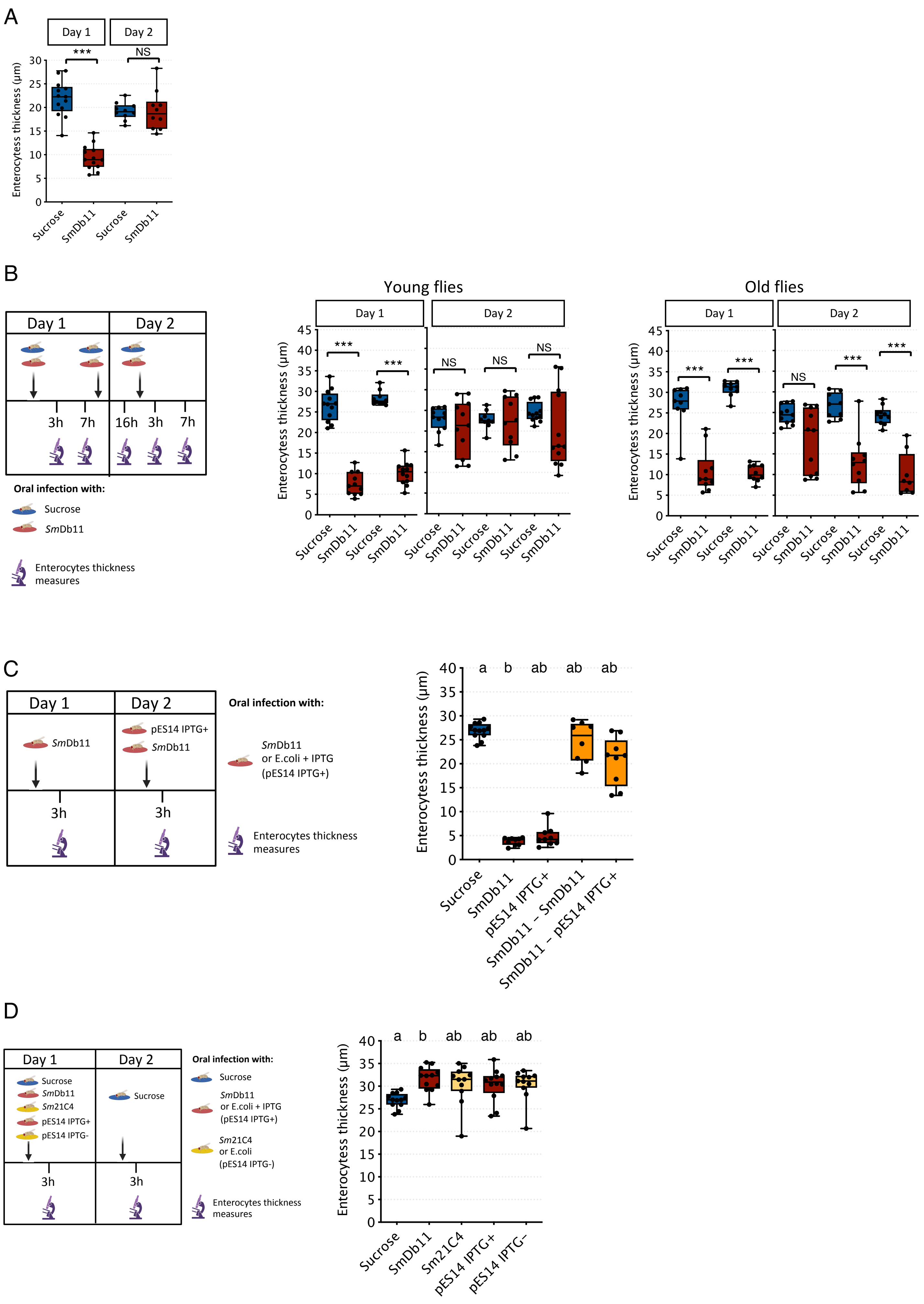
A
Day 1
Day 2
***
NS
Enterocytess thickness (µm)
Sucrose
SmDb11
B
Day 1
Day 2
3h
7h
16h
Oral infection with:
Sucrose
SmDb11
Enterocytes thickness measures
Young flies
Old flies
***
NS
C
SmDb11
pES14 IPTG+
Oral infection with:
SmDb11
or E.coli + IPTG
(pES14 IPTG+)
a
b
ab
pES14 IPTG+
SmDb11 – SmDb11
SmDb11 – pES14 IPTG+
D
Sm21C4
pES14 IPTG-
Sm21C4
or E.coli
(pES14 IPTG-)
pES14 IPTG–

**Fig. S2**

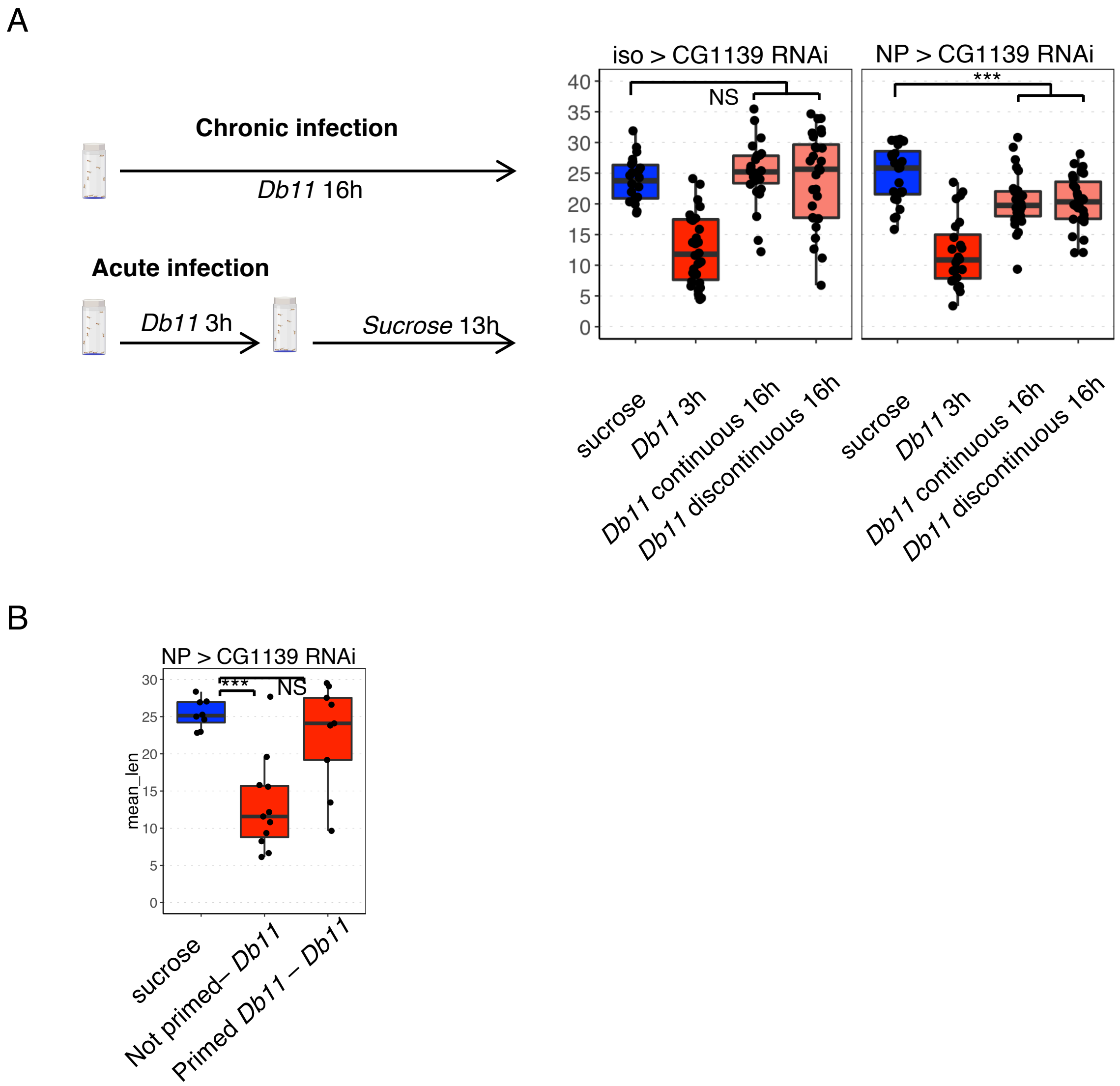
A
Chronic infection
Db11 16h
Acute infection
Db11 3h
Sucrose 13h
iso > CG1139 RNAi
NP > CG1139 RNAi
NS
***
40
35
30
25
20
15
10
5
0
sucrose
Db11 3h
Db11 continuous 16h
Db11 discontinuous 16h
B
NP > CG1139 RNAi
mean_len
***
NS
sucrose
Not primed– Db11
Primed Db11 – Db11

**Fig. S3**
